# *In vivo* imaging and regression analysis reveal cell-non-autonomous functions of Shroom3 during neural tube closure

**DOI:** 10.1101/2022.06.12.495829

**Authors:** Austin T. Baldwin, Juliana H. Kim, John B. Wallingford

## Abstract

During neural tube closure, neural ectoderm cells constrict their apical surfaces to bend and fold the tissue into a tube that will become the central nervous system. These cells are physically interconnected via N-cadherin, and mutation of critical genes within relatively small numbers of cells can result in neural tube closure defects due to non-cell autonomous cell behavior defects. Despite this finding, we have a poor understanding of how neuroepithelial cells interact during apical constriction. In our previous paper, we introduced an imaging and analysis paradigm for tracking and quantifying apical constriction during neural tube closure, while also using CRISPR/Cas9 to generate mosaic loss of function of the apical constriction gene *shroom3*. Here we analyze the behaviors of cells along the mosaic interface of our *shroom3* crispant clones, and find that Shroom3 non-cell autonomously regulates apical constriction and N-cadherin localization. Control cells along the interface constrict less, while *shroom3* crispant cells along the interface constrict more. Finally, we construct a partial least squares regression (PLSR) model to estimate how both autonomous and non-cell autonomous dynamics of actin and N-cadherin affect apical surface area in both control and *shroom3* crispant cells. Overall, our results demonstrate a previously unidentified non-cell autonomous role for Shroom3 in neural tube closure.

## Introduction

Embryonic morphogenesis involves a variety of cellular processes and behaviors that interact to shape a simple cluster of cells into an adult organism. In vertebrate embryos, neural tube closure is the process by which the dorsal neural ectoderm folds itself into a tube that will form the central nervous system. Failure of neural tube closure results in birth defects known as neural tube defects, and outcomes range from serious to deadly (Wallingford et al., 2013). While we possess a wealth of knowledge of genetic factors necessary for neural tube closure(Harris and Juriloff, 2007, 2010), we have a relatively poor understanding of the interplay between various cell behaviors that cooperate to effect successful neural tube closure.

A primary mechanism by which the neural ectoderm folds itself is by apical constriction, where apicobasal polarized cells reduce their apical surface area (Martin and Goldstein, 2014). When neural ectoderm cells fail to apically constrict, neural tube defects such as anencephaly occur (Haigo et al., 2003; Hildebrand and Soriano, 1999; McGreevy et al., 2015). While models of apical constriction typically try to explain the mechanism of constriction within individual cells (Martin and Goldstein, 2014), an epithelium such as the neural ectoderm is composed of numerous cells that are all physically linked to each other by the cadherin/catenin junctional protein complex. Apical constriction depends on actomyosin contractility (Baldwin et al., 2022; Christodoulou and Skourides, 2015; Martin et al., 2009; Roh-Johnson et al., 2012), and apical actomyosin also interacts with the cadherin/catenin complex (Lecuit and Yap, 2015), meaning that actomyosin contractility can be transmitted across entire fields of epithelial cells.

The actin-binding protein Shroom3 is necessary for neural apical constriction and is sufficient to induce apical constriction in apicobasal polarized cells (Haigo et al., 2003; Hildebrand, 2005; Hildebrand and Soriano, 1999). *shroom3* genetically interacts with *N-cadherin* (Plageman et al., 2011), the neural cadherin, and we recently showed that *shroom3* crispant cells are less capable of enriching N-cadherin at both the medial apical surface and apical junctions of neural ectoderm cells (Baldwin et al., 2022). As a classical cadherin, N-cadherin is likely involved in mediating and propagating actomyosin-generated constriction forces across individual apical junctions and subsequently across tissues (Lecuit and Yap, 2015; Nandadasa et al., 2009).

These results suggest the possibility of non-cell autonomous effects, which may impact our understanding of human neural tube defects. Indeed, recent work has found that somatic mutations in several different genes are associated with human NTDs (Tian et al., 2021; Tian et al., 2020) and mosaic alteration of gene function is linked to tissue-wide failure of neural tube closure in both mice and humans (Galea et al., 2021; Kennedy et al., 1998; Rocha et al., 2010). Specifically, mosaic loss of *vangl2* in mice demonstrates that neural tube defects can form in the presence of relatively small numbers of mutant/defective cells (Galea et al., 2021). Because *vangl2* genetically interacts with *shroom3* in the successful completion of neural tube closure/apical constriction (McGreevy et al., 2015), we hypothesize that mosaic loss of *shroom3* may also cause non-autonomous cell behavior defects in the neural ectoderm.

We used our new paradigm of tissue-scale quantification of cell behaviors and protein localization to assess the effect of mosaic loss of *shroom3* in *Xenopus* neural tube closure. Using multivariate regression modeling, we show that medial actin accumulation drives apical constriction non-autonomously in neighborhoods of cells, rather than solely in individual cells. We further show that this neighborhood effect of medial actomyosin contractility on apical surface area is dependent on the function of *shroom3*, both autonomously and non-autonomously. Together these results provide sub-cellular insights into how contractile behaviors within the cells of the neural ectoderm are coordinated and propagated by Shroom3 during neural tube closure.

## Materials and Methods

### Animals

Wild-type Xenopus tropicalis frogs were obtained from the National Xenopus Resource, Woods Hole, MA (Horb et al., 2019).

### Injections

Wild-type X. tropicalis eggs were fertilized in vitro using sperm from wild-type X. tropicalis males using standard methods (Wlizla et al., 2018).

X. tropicalis embryos were moved to 1/9x MMR + 2% Ficoll, then injected in both dorsal blastomeres at the 4-cell stage with 50pg LifeAct-RFP mRNA and 45pg Xenopus N-cadherin-GFP mRNA. In CRISPR-injected embryos, after the next division to reach 8-cell stage, one dorsal blastomere was injected with 1ng Cas9 protein (PNA Bio), 250pg shroom3-targeted sgRNA (target sequence GUAGCCGGAGAGAUCACUUG, Synthego), and 60pg membrane(CAAX)-BFP mRNA. CRISPR validation is described in more detail in (Baldwin et al., 2022).

### Imaging and Cell Segmentation

Injected embryos were held at 25C until they reached Nieuwkoop and Faber (NF) stage 12.5. At NF stage 12.5, vitelline envelopes were removed from embryos and embryos were allowed to “relax” for 30 minutes. Embryos were then mounted in imaging chambers and positioned for imaging of the anterior neural ectoderm. Embryos were imaged on a Nikon A1R confocal microscope using the resonant scanner. Image quality, Z-stacking, and XY tiling were optimized to generate optimal 3D images of the neural plate at a rate of 1 frame per minute. Ultimately, movies of the anterior neural ectoderm of 5 embryos were of sufficient length and quality for analysis; tissue geometry of the initial frame of each of these embryos is presented in (Baldwin et al., 2022).

Beginning at Nieuwkoop & Faber stages 11-12 (Nieuwkoop and Faber, 1994), the anterior neural ectoderm of the injected embryos was subjected to time-lapse confocal microscopy through the process of neural tube closure. Apical cell junctions in each confocal movie were segmented using a combination of Tissue Analyzer, CSML, and EPySeg (Aigouy et al., 2020; Aigouy et al., 2016; Ota et al., 2018). Initial segmentations were hand-corrected using Tissue Analyzer, and then Tissue Analyzer was again used to track cells through each movie.

### Image analysis

Raw 3D images were projected to 2D via maximum intensity and underwent initial segmentation of cell boundaries using the FIJI plugin Tissue Analyzer (Aigouy et al., 2010; Aigouy et al., 2016). The segmentation of an initial frame was hand-corrected, and this hand-corrected segmentation was used to train a classifier using the programs CSML and EPySEG (Aigouy et al., 2020; Ota et al., 2021). CSML and EPySEG were used to generate segmentation for subsequent frames, which were then further hand-corrected in Tissue Analyzer.

After hand-correction, Tissue Analyzer was used to track both cell surfaces and cell junctions, then generate a database of measurements of size and fluorescent intensities for each cell and junction over time. Values for medial and junctional localization of imaged markers in cells were calculated as average pixel fluorescence intensity across the entirety of each respective domain (i.e., total fluorescence of a region divided by the area of the region). Similarly, localization of imaged markers to individual junctions was calculated as an average across the entire junction.

### Data analysis

Cell tracks shorter than 30 frames and junction tracks shorter than 15 frames were discarded.

For individual junctions, errors in junction length caused by Z-displacement and projection were corrected in Matlab.

Cells were determined to be wild-type versus shroom3 crispant by a membrane-BFP localization threshold specific to each embryo. Crispant calls were then manually annotated in cases along the mosaic interface where thresholding produced crispant calls deemed incorrect.

Individual junctions were determined to be wild-type versus shroom3 crispant versus mosaic interface based on the status of the cells the junction was situated between. Wild-type junctions are situated between two wild-type cells, shroom3 crispant junctions are situated between two shroom3 crispant cells, and mosaic interface junctions are situated between a wild-type and a shroom3 crispant cell.

Individual cell and junction tracks were smoothed by averaging over a 7 frame (7 minute) window. Individual cell tracks were further mean-centered and standardized so that variables are measured in standard deviations rather than fluorescence or size units. This standardization allows us to analyze dynamics of cell size and protein localization across a population of cells while controlling for initial size and fluorescence of cells, and was essential for performing partial least squares regression analysis.

Tissue Analyzer databases were imported to R and further analyzed and manipulated primarily using the tidyverse package (R Core Team, 2020; Wickham et al., 2019).

More details on data analysis are included in (Baldwin et al., 2022). In the course of the analysis presented in this manuscript, a small number of errors in cell tracking and *shroom3* crispant calling in (Baldwin et al., 2022) were identified and corrected, so the published dataset in this paper will differ slightly from (Baldwin et al., 2022).

Partial least squares regression models were generated using the R package *“pls”* (Mevik and Wehrens, 2007), using the SIMPLS algorithm (de Jong, 1993). Prior to inclusion in regression models, each variable was mean-centered to zero and scaled by standard deviations within each cell track as previously described in (Baldwin et al., 2022).

## Results

### Heterogeneous apical actin dynamics in the *Xenopus* neural tube

Previously, we performed live imaging and segmentation to capture and quantify cell behavior across large swaths of cells in the closing neural tube. Using this dataset, we described correlations between actin localization and apical surface area in the anterior neural ectoderm. A major conclusion of that study was that apical constriction requires two distinct actin populations, one at the cell-cell junctions and another across the medial apical surface of cells (**Fig. 1A**) (Baldwin et al., 2022). A similar mechanism has been reported in invertebrates (Martin et al., 2009; Roh-Johnson et al., 2012).

**Figure 1:**
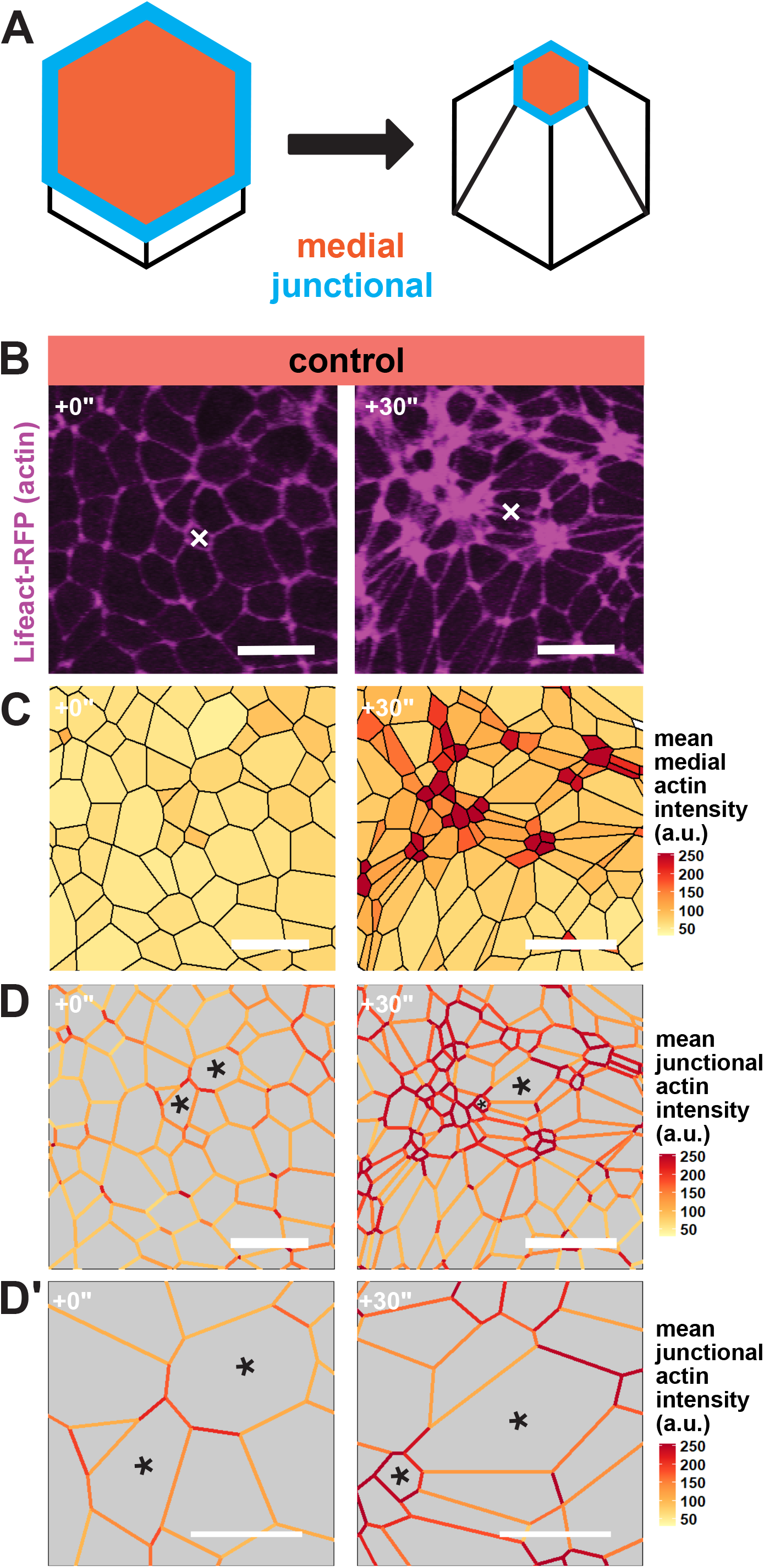
Apical actin accumulation is heterogenous in the *Xenopus* neural ectoderm. **A**, Diagram of medial and junctional quantification domains on the apical surface of apically constricting cells. **B**, Lifeact-RFP localization in the anterior neural ectoderm of apically constricting cells. White “X” marks the same cell in each panel. Scale bar = 25µm. **C**, mean actin intensity at the medial domain of neural ectoderm cells measure in arbitrary units. Scale bar = 25µm. **D**, mean actin intensity at individual cell-cell junctions, measured in arbitrary units. Black asterisks mark the same cells in each frame. Scale bar = 25µm. **D’**, insets of cells from **D**. Scale bar = 12.5µm.

Interestingly, however, when observing changes in actin localization over groups of cells, we noted that cells did not constrict nor accumulate apical actin with perfect uniformity (**Figure 1B**). Rather, it appeared that medial actin contractility was enriched in scattered clusters of cells rather than uniformly across the tissue (**Figure 1B, C**). Additionally, as cells constricted and the tissue deformed, we observed that individual junctions were quite heterogeneous. Indeed, the several cell-cell junctions that bounded each individual cells displayed highly variable actin dynamics. (**Figure 1D, green asterisks**). We provide a higher magnification view of junctional actin intensity in **Figure 1D’** to emphasize the heterogeneous nature of junctional actin dynamics within both individual cells and groups of cells.

### Regression analysis of apical constriction variables reveals non-autonomous effects

Our finding that junctional actin dynamics were heterogeneous suggested that control of actin levels at any given cell-cell junction was not entirely under cell-autonomous control. Given that actomyosin at the apical junctions of epithelial cells is physically linked from cell to cell by cadherins, we hypothesized that our dataset could be used to assess local feedback within groups of cells by calculating the average behavior of both a cell and its neighbors side-by-side (**Figure 2A, 2B**).

**Figure 2:**
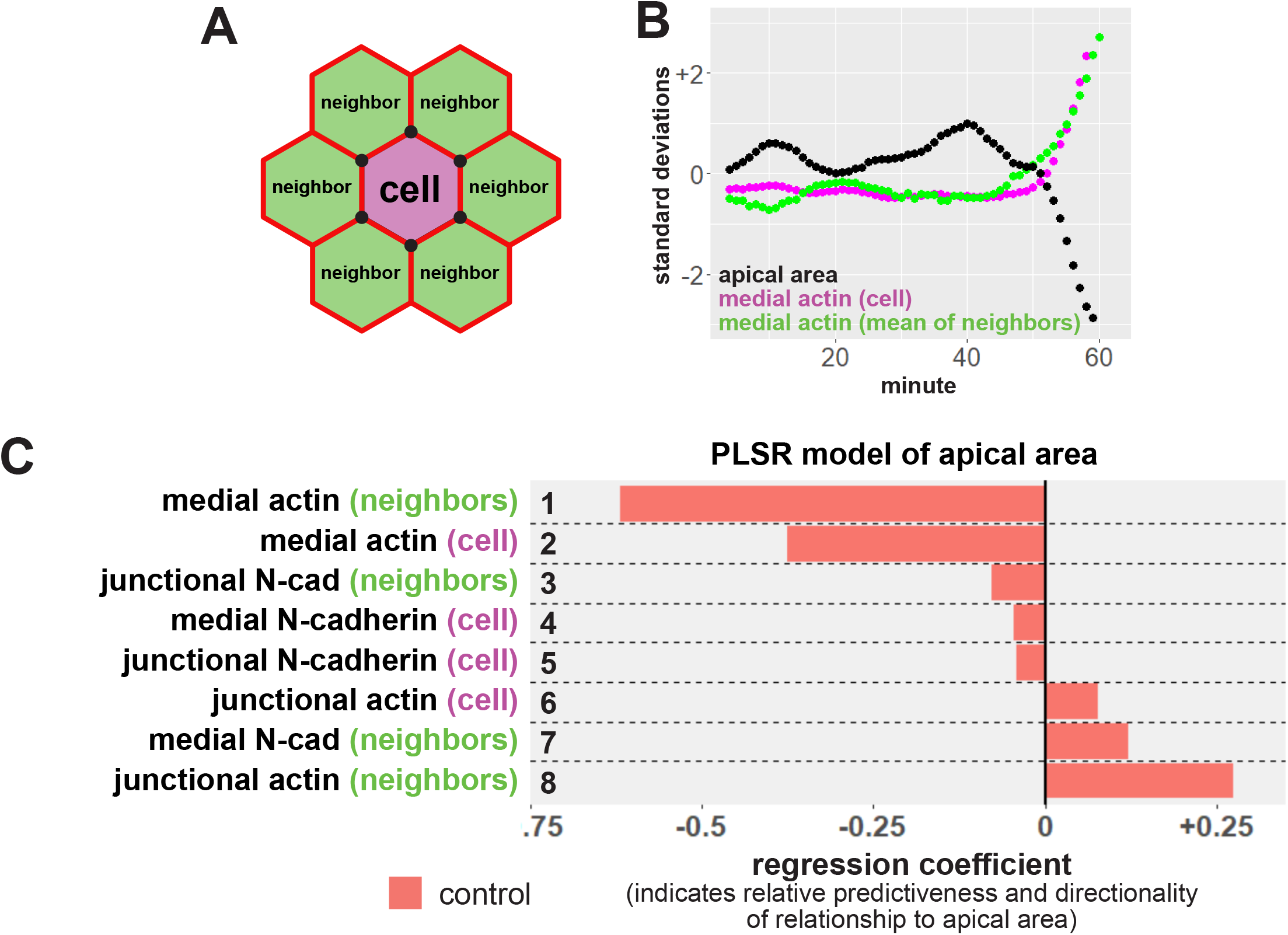
Neighbor analysis and partial least squares regression reveal multiple modes of non-cell autonomous feedback on apical constriction. **A**, diagram of neighbor analyses. Neighbor relationships in each frame are determined by sharing of at least one cell vertex, represented by black circles. Average values for the neighbors of each cell were then calculated. **B**, sample track of a cell and its neighbors. Apical area is shown in black, mean medial actin localization within the cell in magenta, and mean medial actin of neighboring cells shown in green. s.d. = standard deviations. **C**, partial least squares regression (PLSR) of apical area in control cells in the anterior neural ectoderm. Regression coefficients represent the independent effect of each variable on apical surface area.

Our analysis pipeline collects a number of measurements of cell size and protein localization for each observation, and by coupling the quantification of each cell’s behavior to the mean of all of that cell’s neighbors, we introduce tremendous complexity. To address this large degree of complexity in an unbiased way, we examined relationships between variables using Partial Least Squares Regression (PLSR). This multivariate regression technique predicts the behavior of a single response variable based on the behavior of multiple other, potentially-covarying, test variables in the dataset (e.g., (Janes et al., 2005; Janes and Yaffe, 2006). PLSR models provide regression coefficients that reflect the relative magnitude of the independent effect of each test variable on the response variable (Mevik and Wehrens, 2007; Palermo et al., 2009). Specifically, we used PLSR modeling to ask how the change in a cell’s apical area (response variable) was predicted by the dynamics of actin and N-cadherin localization in a given cell and in that cell’s neighbors (test variables).

Our initial regression model was generated from ∼65,000 observations of ∼1,100 control cells (from 5 embryos), and immediately yielded interesting results (**Figure 2C**). First, and to our surprise, by far the strongest predictor of apical area was the intensity of medial actin *in neighboring cells* (**Figure 2C, row 1**). Medial actin within the cell was the second most predictive factor, but was less predictive than neighbor’s medial actin (**Figure 2C, row 2**). Both of these variables were negative predictors of area (i.e. as actin localization increases, apical area decreases). These results agree with our previous data demonstrating that medial actin localization is better correlated with apical constriction than is junctional actin (Baldwin et al., 2022) and moreover suggests that the medial actin network exerts *non-autonomous* effects on apical constriction in the neural epithelium.

The second surprising result from our PLSR was that junctional actin intensity in neighboring cells emerged as a *positive* predictor of a cell’s apical area (**Figure 2C, row 8**), suggesting that increasing junctional actin in a cell’s neighbors will inhibit that cell’s apical constriction. Thus, multiple influential components in our PLSR model paint a picture of a key role for cell non-autonomous effects in apical constriction in the closing *Xenopus* neural tube.

### Mosaic junctions linking control and *shroom3* crispant cells display a distinct phenotype

Previously, we described generation of embryos with mosaic loss-of-function of the gene *shroom3* via CRISPR/Cas9 microinjection (Baldwin et al., 2022). In that work, we focused on cells and junctions fully within control or crispant clones (i.e. considering only cells surrounded by like cells). However, those datasets also contained many cells that lay on the interface between those clones (**Figure 3A**), and our tracking and analysis methodology allowed us to automatically identify junctions along the mosaic interface between clones and analyze them as a distinct category (**Figure 3B**). Using this approach, we found that mosaic interface junctions presented an intermediate constriction phenotype, shrinking significantly less than junctions linking two control cells and significantly more than junctions linking two *shroom3* crispant cells (**Figure 3C**).

**Figure 3:**
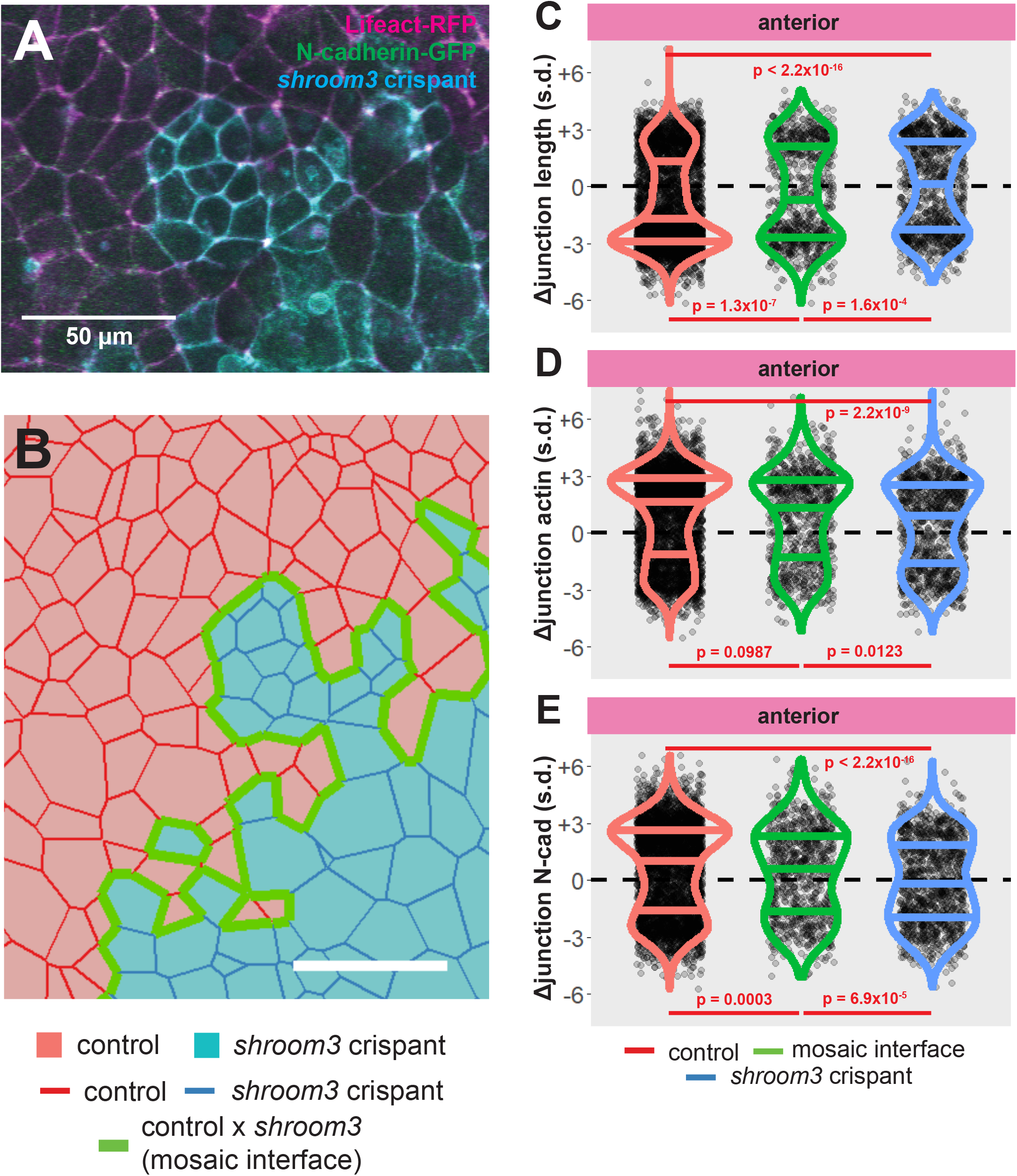
Loss of Shroom3 disrupts junction behaviors along the mosaic interface. **A**, control and *shroom3* crispant cells meet at the mosaic interface. Magenta = Lifeact-RFP/actin, green = N-cadherin-GFP, cyan = membrane(CAAX)-BFP + Cas9 protein (PNA Bio) + *shroom3-*targeted sgRNA. Scale bar = 50µm. **B**, cell tracking can automatically identify cell junctions at the mosaic interface. Red cells = control cells, blue cells = *shroom3* crispant cells. Red junctions = control x control interaction, green junctions = control x *shroom3* crispant interaction (mosaic interface), blue junctions = *shroom3* crispant x *shroom3* crispant interactions. Scale bar = 50µm. **C, D, E**, overall change in junction length, mean junctional actin, and mean junctional N-cadherin in control, mosaic interface, and *shroom3* crispant junctions. Each dot is an individual junction. Horizontal lines within each violin delineate quartiles along each distribution. s.d. = standard deviations. P-values were calculated via KS test.

To understand the origin of this phenotype, we examined the dynamics of actin and N-cadherin, both of which increase significantly at cell-cell junction in normal cells (**Fig. 3D, E, red violins**; see also (Baldwin et al., 2022)). We found no difference in actin accumulation between control and mosaic interfaces (**Fig. 3D, green violin**), despite their highly significant defect in constriction (**Fig. 3C, green violin**). By contrast, accumulation of N-cadherin along mosaic interface junctions was significantly disrupted compared to control junctions (**Fig. 3E, green violin**). Thus, defective constriction at interface junctions joining control and *shroom3* crispant cells does not arise from a failure of actin assembly at the junction but may reflect in part a relate to defects in junctional N-cadherin.

### Control cells at the interface of *shroom3* crispant clones display non-autonomous apical constriction defects

In addition to identifying junctions between control and *shroom3* crispant cells, our analysis platform allows us to track the behavior of cells located at the mosaic interface. For brevity, cells located completely within a clone will be labelled “clone” (i.e. control cells apposed only to control cells, or *shroom3* cells apposed only to *shroom3* cells), and cells located at the mosaic interface will be labeled “interface” (i.e. control cells apposed to *shroom3* crispant cells) (**Figure 4A**).

**Figure 4:**
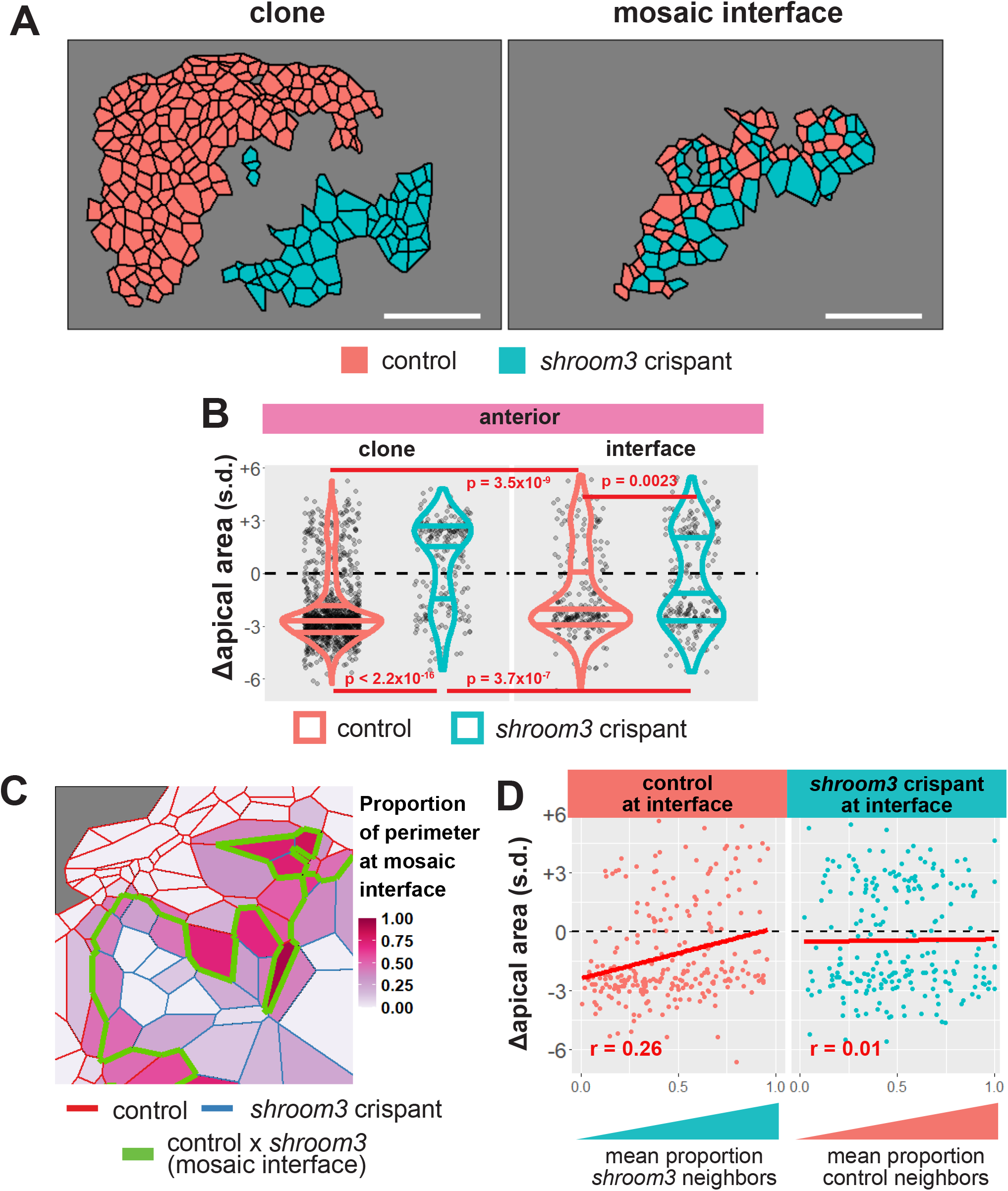
Loss of Shroom3 causes non-cell autonomous apical constriction defects along the mosaic interface. **A**, sample embryo showing cells within control and *shroom3* crispant clones (left panel) and cells located at the mosaic interface (right panel). Red = control cells, blue = *shroom3* crispant cells. **B**, change in apical area among control and *shroom3* crispant cells both within clones (left) and at the mosaic interface (right). Red violins = control cells, blue violins = *shroom3* crispant cells. Each dot is an individual cell. Horizontal lines within each violin delineate quartiles along each distribution. s.d. = standard deviations. P-values were calculated via KS test. **C**, schematic of calculation of proportion of neighbors at mosaic interface. **D**, change in apical area versus proportion of cell boundary with unlike neighbors (i.e. at mosaic interface) for both control and *shroom3* crispant cells at the mosaic interface. s.d. = standard deviations. R-values calculated by Pearson’s correlation.

As previously reported in (Baldwin et al., 2022), *shroom3* crispant clone cells display significantly reduced apical constriction compared to control clone cells, and most *shroom3* clone cells actually dilate their apical surfaces over time (**Figure 4B, left**). Interestingly, however, controls cells at the interface also display a significant apical constriction defect as compare to control clone cells (**Figure 4B**). Conversely, *shroom3* crispant cells at the interface constrict significantly *more* than do *shroom3* crispant clone cells (**Figure 4B**). Thus, both control and *shroom3* crispant cells display non-autonomous alterations to apical constriction when present at a mosaic interface.

To explore this effect further, we calculated the proportion of the perimeter of each interface cell that lies along the mosaic interface (i.e. the proportion of perimeter that a control cell shares with *shroom3* crispant cells, or vice versa) (**Figure 4C**). For each cell, we calculated the average proportion of the cell’s perimeter that was at the interface over the tracked period and compared that to the amount of apical constriction that cell achieved.

In control cells at the interface, we found that increasing amounts of mosaic interface correlated with increasingly defective apical constriction (r = 0.26, Pearson’s correlation), such that control cells were less able to reduce apical area as they became more surrounded by *shroom3* cells (**Figure 4D, left panel**). This result suggests that Shroom3 is required in the neighboring cell to exert the normal non-cell autonomous effect to promote apical constriction that we predicted by our PLSR model (**Figure 2C, Line 1**). Conversely, *shroom3* crispant cells at the interface showed essentially no correlation between their proportion of control neighbors and the amount of apical constriction they achieved (**Figure 4D, right panel**). This result, in turn, suggests that Shroom3 is also required in a constricting cell to receive or respond to the non-autonomous effect exerted by neighboring control cells.

### The non-autonomous effect of medial actin accumulation on apical constriction is disrupted between control and *shroom3* crispant cells

Our data thus far suggest that Shroom3 is required both in apically constricting cells and in their neighbors to effect normal apical constriction, so we turned again to PLSR for insight. As we did for control cells in Figure 2, we calculated neighbor means for *interface* cells, but we separately calculated averages of control neighbors and *shroom3* crispant neighbors for each cell (**Figure 5A, B**). This dataset was then used to calculate new PLSR models, one for apical area in control cells at the mosaic interface and another for *shroom3* crispant cells at the interface.

**Figure 5:**
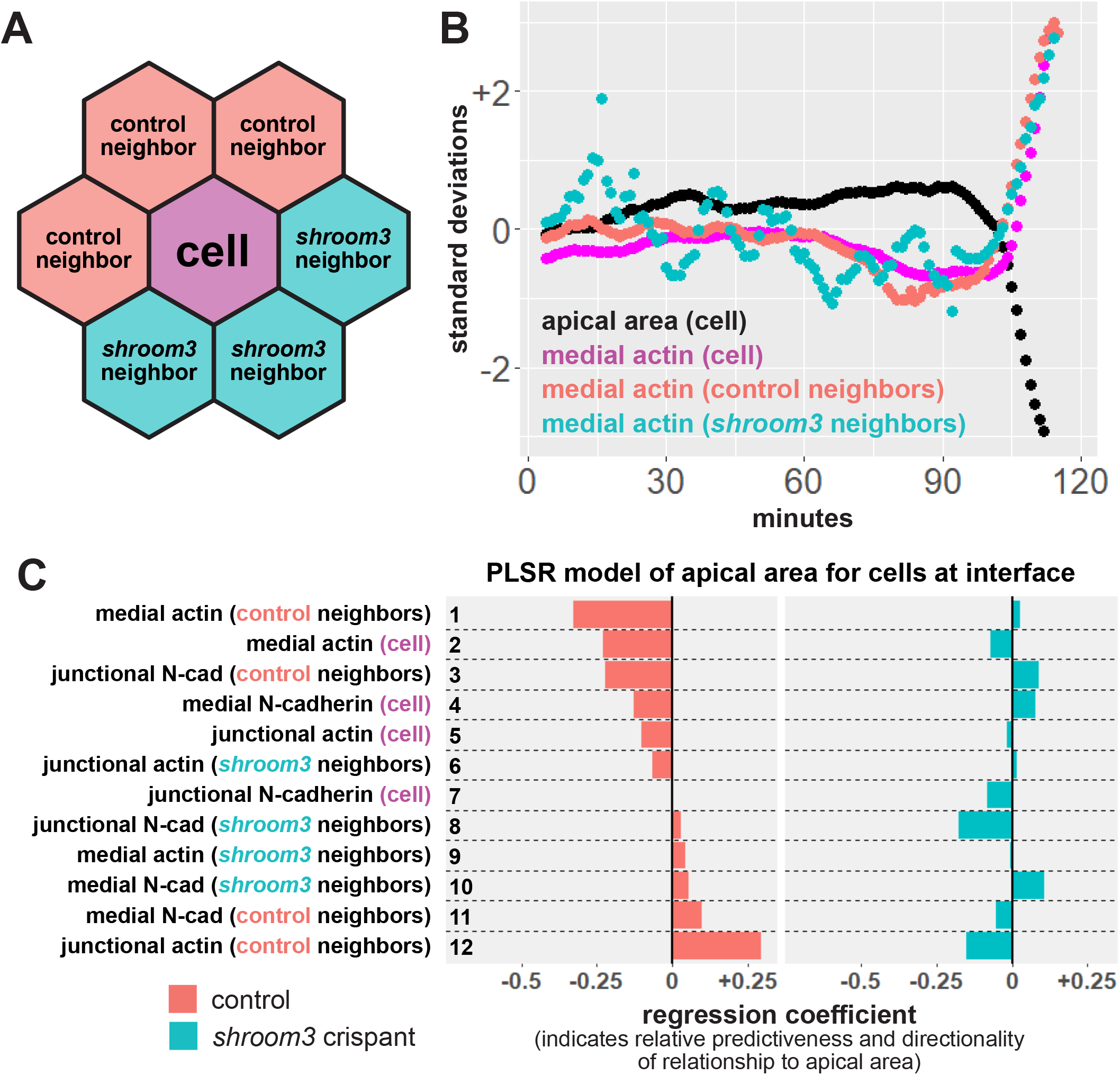
Loss of Shroom3 disrupts non-cell autonomous effects of actin localization. **A**, schematic of neighbor analysis of cells along the mosaic interface. Mean behaviors of both control and *shroom3* crispant neighbors can be calculated separately. **B**, sample cell track showing the behavior of one control cell at the mosaic interface and its neighbors. Black = apical area of cell, magenta = mean medial actin of cell, red = mean medial actin of control neighbors, blue = mean medial actin of *shroom3* crispant neighbors. **C**, partial least squares regression (PLSR) of apical area in both control and *shroom3* crispant cells along the mosaic interface. Protein localization variables on the right indicate whether the resulting coefficient was generated based on influence of the variable within a cell, from a cell’s control neighbors, or a cell’s *shroom3* crispant neighbors. Regression coefficients show the influence of each variable on apical area in both control cells (red bars) and *shroom3* crispant cells (blue bars).

In control cells, we found that the strongest predictors of apical area were medial actin localization of the neighbors and of the cell itself (**Figure 5C, red bars, rows 1 & 2**), similar if somewhat weaker than we observed in the model of clone control cells (see **Figure 2C, rows 1 & 2**). Strikingly, however, in *shroom3* crispant cells, medial actin of neither neighbors nor of the cell itself was predictive of apical area (**Figure 5C, blue bars, rows 1 & 2**). The latter result is consistent with our previous finding that medial actin localization is poorly correlated with apical area in *shroom3* cells (Baldwin et al., 2022), and both results are consistent with data in Figure 4, suggesting that loss of *shroom3* bi-directionally disrupts the non-cell autonomous effect of medial actin accumulation to induce apical constriction across groups of cells.

## Conclusions

So far, relatively little is known about non-autonomous functions of “morphogenesis genes” such as *shroom3*. That said, mosaic disruption of a variety of signaling, adhesion, and contractility genes in *Drosophila* and the planar polarity gene *Dchs1* in mice have been shown to alter behaviors and protein localization in neighboring cells (Chan et al., 2017; Franke et al., 2005; James et al., 2002; Laplante and Nilson, 2006). Indeed, mosaic loss of *N-cadherin* is sufficient to disrupt tissue structure in both *Drosophila* and zebrafish, again highlighting the importance of the interactions of individual cell to overall tissue morphogenesis (Chan et al., 2017; McMillen et al., 2016). Most recently, a paper by Galea *et al. (Galea et al*., *2021)* analyzed non-autonomous defects of apical constriction using mosaic loss-of-function of the Planar Cell Polarity (PCP) gene *vangl2*; given the evolutionarily conserved non-autonomy of PCP signaling (Adler et al., 2000; Phillips et al., 2008), non-autonomous effects on cell behaviors were fascinating but not perhaps not surprising. Conversely, Shroom3 is both necessary and sufficient to induce apical constriction in apicobasal polarized cells (Haigo et al., 2003), but to date there has been little evidence to suggest it may act non-autonomously.

Here, using tissue scale imaging of cell behaviors, unbiased regression analysis of the data, and mosaic knockdown, we show that *shroom3* exerts both cell-autonomous and cell-non-autonomous effects on apical constriction in the closing *Xenopus* neural tube. Our data suggest the presence of an as yet undefined, Shroom3-dependent positive feedback loop that links medial actin dynamics and apical constriction. Data here and in our previous paper (Baldwin et al., 2022) suggest that N-cadherin also plays a key role here. Future work should focus on higher image and time resolution observations of the interactions between control and *shroom3* loss-of-function cells, and detailed analysis of the non-autonomous functions of other neural tube closure genes will also be of great interest.

## Acknowledgements

We thank Robert Huebner, Elle Roberson, Caitlin Devitt, and Haleigh Mendiola for critical reading. This work was supported by the NICHD, including R01HD099191 to JBW and F32HD094521 to A.B.

